# Biosynthesis of lavandulol and lavandulyl acetate in *Escherichia coli*

**DOI:** 10.1101/2025.06.28.662091

**Authors:** Dianqi Yang, Xiaoqiang Ma

## Abstract

Lavandulol and its ester derivative, lavandulyl acetate, are key fragrance constituents of lavender essential oil with widespread applications in cosmetics, perfumery, and food. However, traditional plant extraction suffers from low yield and unsustainable practices, and chemical synthesis relies on petrochemical feedstocks. Here, we report the first *de novo* microbial biosynthesis of lavandulol and lavandulyl acetate in *Escherichia coli* through modular pathway engineering. By screening 13 pyrophosphatases from *E. coli*, we identified RdgB as an efficient pyrophosphatase catalyzing the conversion of lavandulyl diphosphate into lavandulol, enabling its production at 24.9 mg/L. Building on this, we established a three-plasmid expression system and introduced a lavender-derived alcohol acyltransferase, LiAAT4, to achieve the biosynthesis of lavandulyl acetate at 42.4 mg/L. This study establishes the biosynthetic route to lavandulol in *E. coli*, demonstrating a viable and sustainable microbial platform for producing significant natural components of lavender oil. Our work provides the basis for future strain optimization and industrial-scale biomanufacturing of high-value monoterpene fragrance molecules.

## 1. Introduction

Monoterpenes represent a diverse class of terpenoid natural products derived from the condensation of two five-carbon isoprene units[1], widely distributed in plants, insects, and microbes, and extensively utilized in fragrances, food additives, and other applications[2, 3]. Structurally, monoterpenes are categorized as either regular that is biosynthesized from the head-to-tail condensation of dimethylallyl diphosphate (DMAPP) and isopentenyl diphosphate (IPP) or irregular, which arise through alternative linkage patterns such as head-to-middle condensation of two DMAPP units[4]. Lavandulol is a representative irregular monoterpene alcohol that constitutes one of the major active components of lavender essential oil. It contributes significantly to the oil’s characteristic floral aroma and exhibits bioactive properties that underpin its use in perfumery, food flavoring, and cosmetic formulations[5, 6]. As demand for natural and sustainable fragrance molecules continues to rise, lavandulol and its derivatives are increasingly recognized as high-value targets for industrial biosynthesis.

Currently, the commercial supply of lavandulol relies on direct extraction from lavender plants[7]. However, this approach suffers from several inherent limitations. The agricultural production of lavender is constrained by long growth cycles, seasonal variation, and labor-intensive harvesting, which collectively contribute to high costs and inconsistent yields. Moreover, the lavandulol content in ethanol extracts from lavender flowers is relatively low, typically around 1.54 % (w/w)[8], making extraction-based approaches impractical for meeting large-scale industrial demand. Besides, chemical synthesis routes depend on non-renewable petrochemical feedstocks and often involve environmentally burdensome processes. These limitations underscore the urgent need for alternative production platforms that are scalable, sustainable, and economically viable.

Microbial cell factories offer a promising solution through the integration of synthetic biology and metabolic engineering. These platforms enable the conversion of simple carbon sources into complex natural products via rationally designed biosynthetic pathways. Microbial biosynthesis provides key advantages over plant extraction and chemical synthesis, including reduced dependence on land and climate, simplified processing steps, shorter production cycles, and avoidance of hazardous reagents or waste byproducts. Despite these benefits, the microbial biosynthesis of lavandulol has remained largely unexplored due to an incomplete understanding of its biosynthetic pathway. Recent progress by Nie et al.[9] demonstrated the feasibility of *de novo* lavandulol biosynthesis in *Saccharomyces cerevisiae* by functionally expression and protein engineering of a lavandulyl diphosphate synthase from *Lavandula x Intermedia* (LiLPPS). Through combinatorial pathway engineering, including overexpression of the genes coding the rate-limiting enzymes (*IDI1*, *tHMG1*), gene deletions (*MLS1*, *CIT2*), and promoter replacements, they achieved lavandulol titers of up to 308.92 mg/L in a fed-batch system. However, the absence of a complete, modular microbial chassis and the lack of alternative esterification biosynthetic routes using available alcohol acetyltransferase (AAT) still limit the broader application of its ester derivatives.

In this study, we addressed these challenges by systematically exploring the biosynthetic potential of *Escherichia coli* for lavandulol production. We performed a comprehensive screening of 13 endogenous pyrophosphatases (PPase) and identified RdgB as a previously unrecognized yet highly efficient phosphatase for converting lavandulyl diphosphate (LPP) to lavandulol via the mevalonate (MVA) pathway with Ethanol Utilization Pathway (EUP)[10] to supply more primer substrate. This finding enabled the construction of a robust *E. coli* biosynthetic chassis capable of lavandulol production. To further expand the product spectrum, we introduced LiAAT4, an AAT from *Lavandula* x *intermedia*[11], under a cumate-inducible promoter, enabling the *in vivo* esterification of lavandulol to lavandulyl acetate, another industrially valuable derivative. The resulting three-plasmid expression system achieved lavandulol and lavandulyl acetate titers of 24.9 mg/L and 42.4 mg/L, respectively, under shake-flask conditions with *in situ* extraction. This study represents the successful reconstruction of the complete microbial biosynthetic route to lavandulol and its acetate in *E. coli*, demonstrating both enzymatic feasibility and pathway modularity. These findings not only provide fundamental insights into irregular monoterpene biosynthesis but also establish a platform for further optimization toward biomanufacturing of high-value fragrance molecules.

## 2. Materials and Methods

### 2.1 Chemical, strain and medium

Racemic lavandulol (BCRYV130780) was purchased from BOCSCI Inc., racemic lavandulyl acetate (HY-117419A) was purchased from MedChemExpress (MCE, Monmouth Junction, NJ, USA). Tryptone (LP0042B) and yeast extract (LP0021B) were purchased from Oxoid (Thermo Fisher Scientific Inc., USA), and all the other chemicals were purchased from Shanghai Aladdin Biochemical Technology Co., Ltd. (Shanghai, China) unless specifically mentioned. *Escherichia coli* DH5α used for cloning and plasmid construction was purchased from Tsingke Biotechnology Co., Ltd. (Beijing, China). Lysogeny Broth (LB) medium composed of 10 g/L tryptone, 5 g/L yeast extract, and 5 g/L NaCl, and autoclaved at 121 °C for 20 min.

*E. coli MG1655 DE3 ΔendA ΔrecA* (https://www.addgene.org/37854/) was purchased from Addgene. The engineered *E. coli* DH5α carrying the recombinant plasmids were usually cultured in M2 medium at 30 °C after adding the inducers. The M2 medium used for lavandulol biosynthesis contained 10 g/L glycerol, 10 g/L tryptone, 5 g/L yeast extract, 13.3 g/L KH_2_PO_4_, and 4 g/L (NH_4_)_2_HPO_4_, and the pH was adjusted to 7.0 with NaOH and autoclaved at 121 °C for 20 min. The TB medium used for lavandulyl acetate biosynthesis contained 10 g/L glycerol, 12 g/L tryptone, 24 g/L yeast extract, 2.2 g/L KH_2_PO_4_, and 9.4 g/L (NH_4_)_2_HPO_4._

### 2.2 Plasmid and strain construction

The genes coding 13 PPases were amplified from the colony PCR using *E. coli MG1655 DE3 ΔendA ΔrecA*. The construction of the plasmids in this study was based on the GT DNA assembly standard[12]. All the plasmids, strains and genes constructed and used in this study are listed in **Table 1-3**. The standard electroporation protocol was used to transform the plasmid into *E. coli*.

**Table 1.** Plasmid information.

| **Name** | **Genotype of plasmids** | **Purpose in this study** |
| --- | --- | --- |
| pMVA | SpecR-pAC-Lacl-pT7_RBS1_EcatoB_RBS2stop_Sahmgs_RBS4stop_Sahmgr-rrnBT-ptrc_RBS1_Scmk_RBS2stop_Scpmk_RBS4stop_Scpmd_RBS3stop_Ecidi-T7T | IPP and DMAPP production from glycerol |
| pLav1+EUP | AmpR-pMB1-Lacl-pT7_RBS1_trLiLPPS-T7T-pLuxB_RBS1_EcNudJ-L3S2P21T-LuxR-pGyrA_RBS1_DpADA_RBS2stop_ScADH2-T7T | Lavandulol production |
| pLav2+EUP | AmpR-pMB1-Lacl-pT7_RBS1_trLiLPPS-T7T-pLuxB_RBS1_EcCdh-L3S2P21T-LuxR-pGyrA_RBS1_DpADA_RBS2stop_ScADH2-T7T | Lavandulol production |
| pLav3+EUP | AmpR-pMB1-Lacl-pT7_RBS1_trLiLPPS-T7T-pLuxB_RBS1_EcPhoA-L3S2P21T-LuxR-pGyrA_RBS1_DpADA_RBS2stop_ScADH2-T7T | Lavandulol production |
| pLav4+EUP | AmpR-pMB1-Lacl-pT7_RBS1_trLiLPPS-T7T-pLuxB_RBS1_EcPpa-L3S2P21T-LuxR-pGyrA_RBS1_DpADA_RBS2stop_ScADH2-T7T | Lavandulol production |
| pLav5+EUP | AmpR-pMB1-Lacl-pT7_RBS1_trLiLPPS-T7T-pLuxB_RBS1_EcRdgB-L3S2P21T-LuxR-pGyrA_RBS1_DpADA_RBS2stop_ScADH2-T7T | Lavandulol production |
| pLav6+EUP | AmpR-pMB1-Lacl-pT7_RBS1_trLiLPPS-T7T-pLuxB_RBS1_EclpxH-L3S2P21T-LuxR-pGyrA_RBS1_DpADA_RBS2stop_ScADH2-T7T | Lavandulol production |
| pLav7+EUP | AmpR-pMB1-Lacl-pT7_RBS1_trLiLPPS-T7T-pLuxB_RBS1_EcMutT-L3S2P21T-LuxR-pGyrA_RBS1_DpADA_RBS2stop_ScADH2-T7T | Lavandulol production |
| pLav8+EUP | AmpR-pMB1-Lacl-pT7_RBS1_trLiLPPS-T7T-pLuxB_RBS1_EcNudC-L3S2P21T-LuxR-pGyrA_RBS1_DpADA_RBS2stop_ScADH2-T7T | Lavandulol production |
| pLav9+EUP | AmpR-pMB1-Lacl-pT7_RBS1_trLiLPPS-T7T-pLuxB_RBS1_EcYtjC-L3S2P21T-LuxR-pGyrA_RBS1_DpADA_RBS2stop_ScADH2-T7T | Lavandulol production |
| pLav10+EUP | AmpR-pMB1-Lacl-pT7_RBS1_trLiLPPS-T7T-pLuxB_RBS1_EcyceF-L3S2P21T-LuxR-pGyrA_RBS1_DpADA_RBS2stop_ScADH2-T7T | Lavandulol production |
| pLav11+EUP | AmpR-pMB1-Lacl-pT7_RBS1_trLiLPPS-T7T-pLuxB_RBS1_Ecppx-L3S2P21T-LuxR-pGyrA_RBS1_DpADA_RBS2stop_ScADH2-T7T | Lavandulol production |
| pLav12+EUP | AmpR-pMB1-Lacl-pT7_RBS1_trLiLPPS-T7T-pLuxB_RBS1_EcnudF-L3S2P21T-LuxR-pGyrA_RBS1_DpADA_RBS2stop_ScADH2-T7T | Lavandulol production |
| pLav13+EUP | AmpR-pMB1-Lacl-pT7_RBS1_trLiLPPS-T7T-pLuxB_RBS1_Ecdut-L3S2P21T-LuxR-pGyrA_RBS1_DpADA_RBS2stop_ScADH2-T7T | Lavandulol production |
| pLiAAT4K | KanR-pCDF-pCym_RBS1_LiAAT4-L3S2P21T-CymR | AAT for esterification |

**Table 2.**
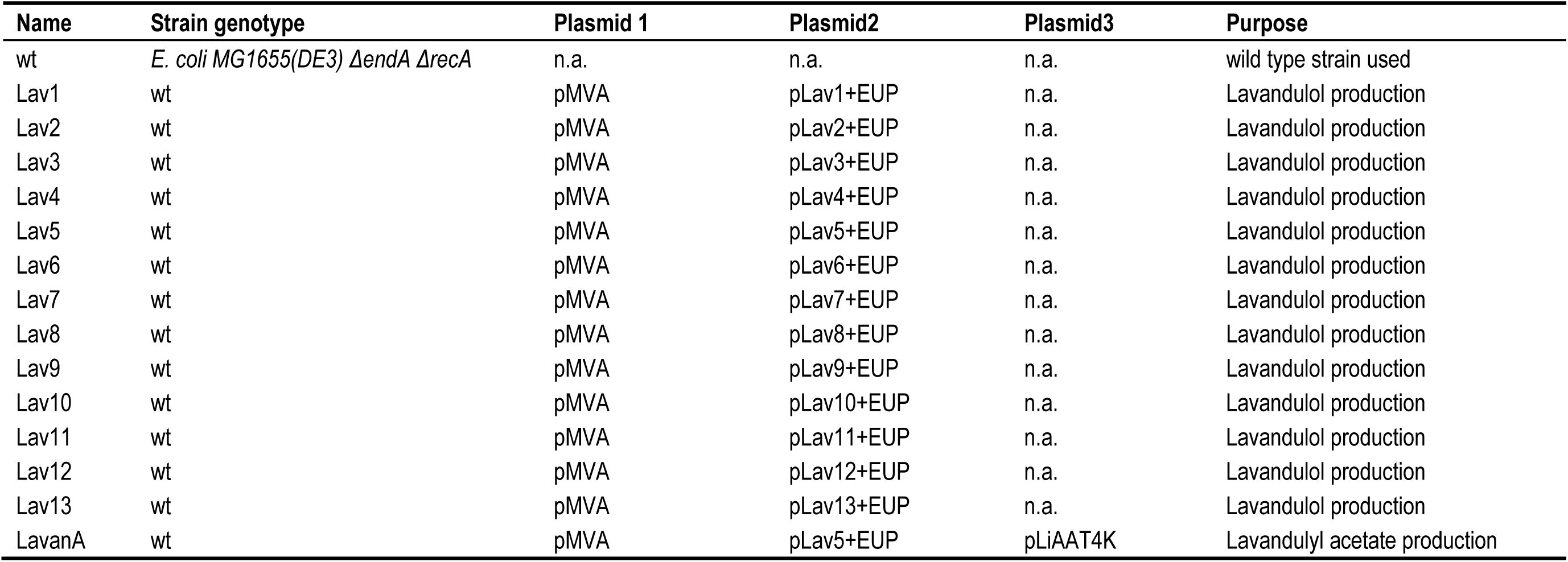
Strain information.

| **Name** | **Strain genotype** | **Plasmid 1** | **Plasmid2** | **Plasmid3** | **Purpose** |
| --- | --- | --- | --- | --- | --- |
| wt | *E. coli* MG1655(DE3) $\Delta endA \Delta recA$ | n.a. | n.a. | n.a. | wild type strain used |
| Lav1 | wt | pMVA | pLav1+EUP | n.a. | Lavandulol production |
| Lav2 | wt | pMVA | pLav2+EUP | n.a. | Lavandulol production |
| Lav3 | wt | pMVA | pLav3+EUP | n.a. | Lavandulol production |
| Lav4 | wt | pMVA | pLav4+EUP | n.a. | Lavandulol production |
| Lav5 | wt | pMVA | pLav5+EUP | n.a. | Lavandulol production |
| Lav6 | wt | pMVA | pLav6+EUP | n.a. | Lavandulol production |
| Lav7 | wt | pMVA | pLav7+EUP | n.a. | Lavandulol production |
| Lav8 | wt | pMVA | pLav8+EUP | n.a. | Lavandulol production |
| Lav9 | wt | pMVA | pLav9+EUP | n.a. | Lavandulol production |
| Lav10 | wt | pMVA | pLav10+EUP | n.a. | Lavandulol production |
| Lav11 | wt | pMVA | pLav11+EUP | n.a. | Lavandulol production |
| Lav12 | wt | pMVA | pLav12+EUP | n.a. | Lavandulol production |
| Lav13 | wt | pMVA | pLav13+EUP | n.a. | Lavandulol production |
| LavanA | wt | pMVA | pLav5+EUP | pLiAAT4K | Lavandulyl acetate production |

**Table 3.**
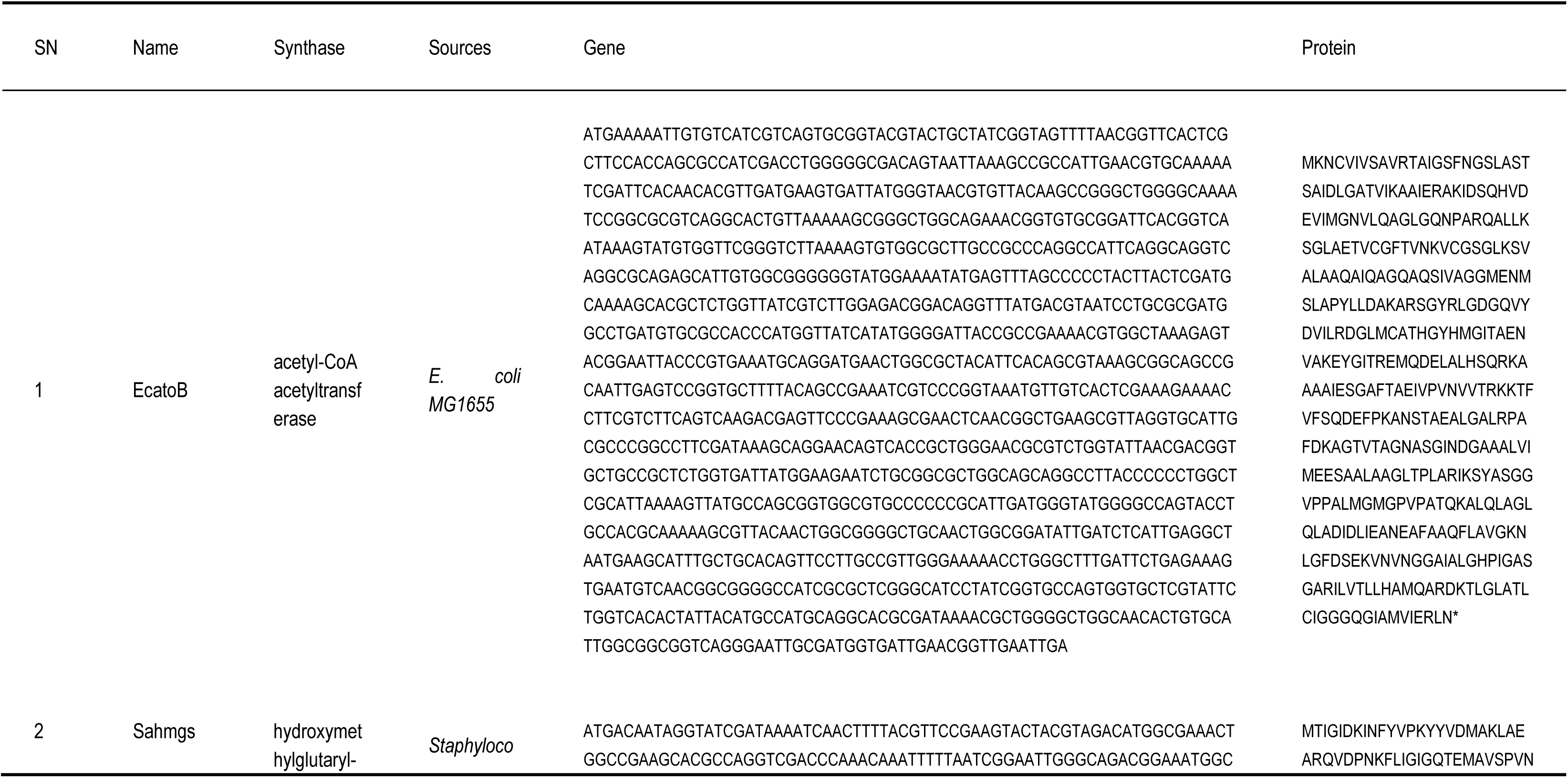
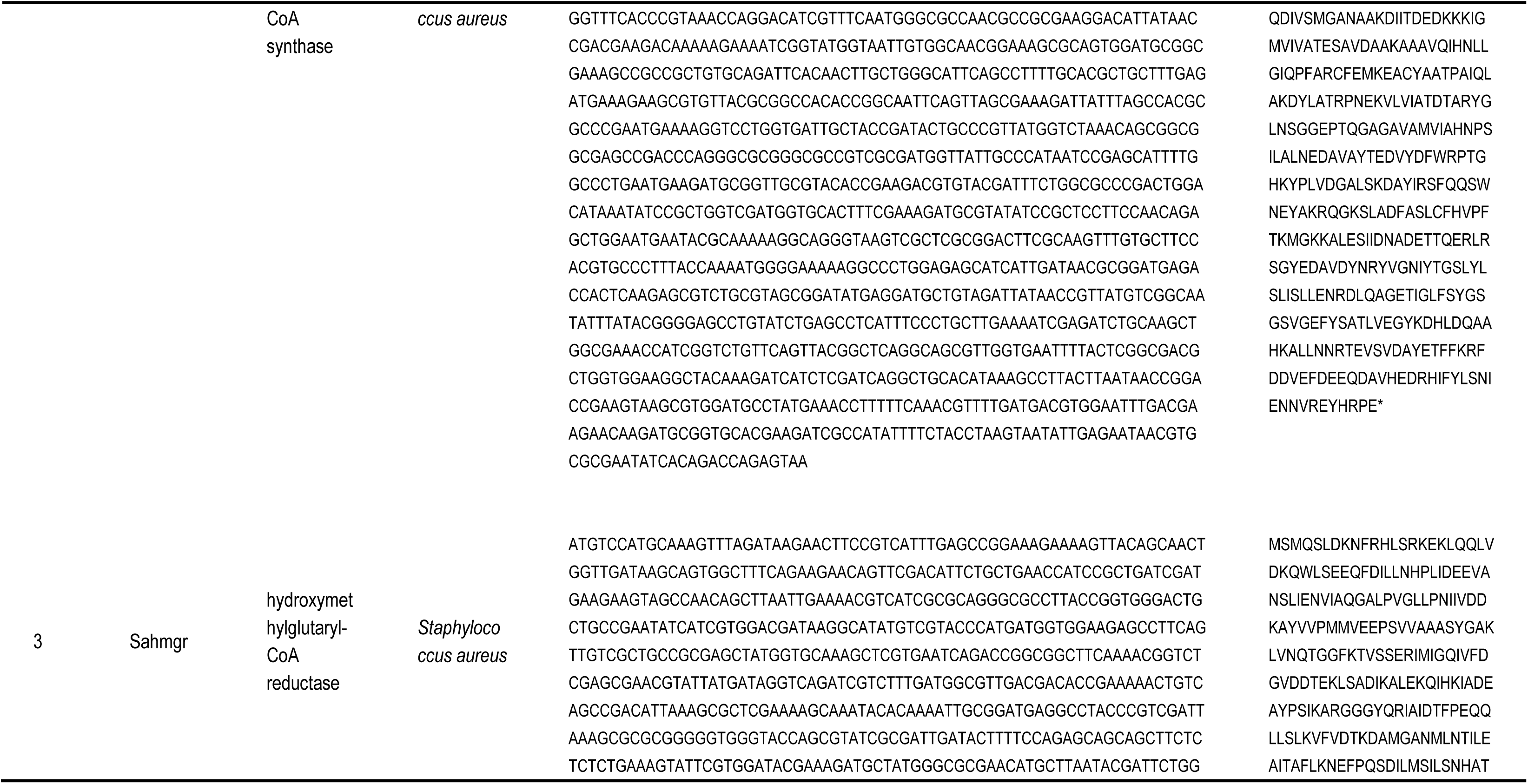
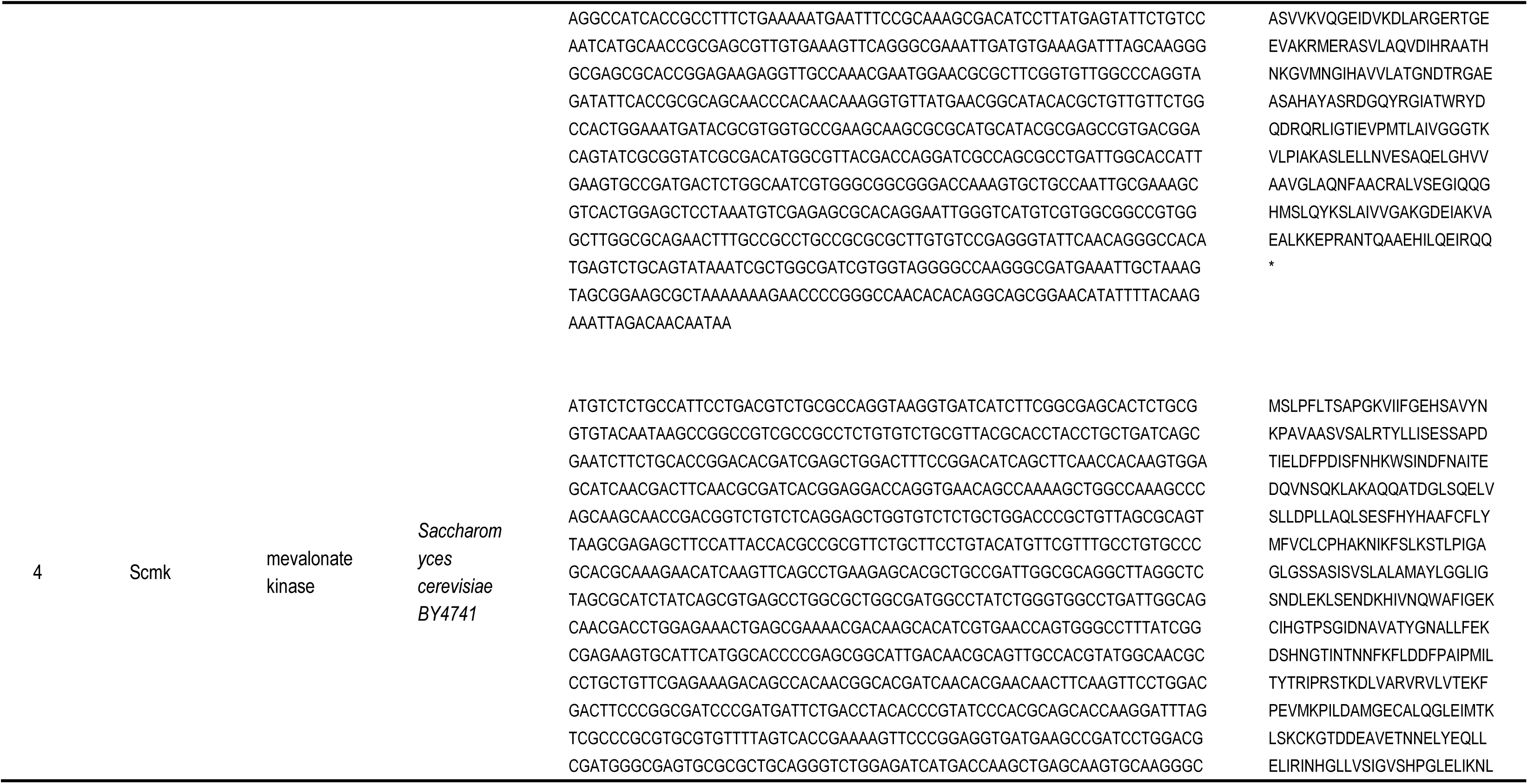
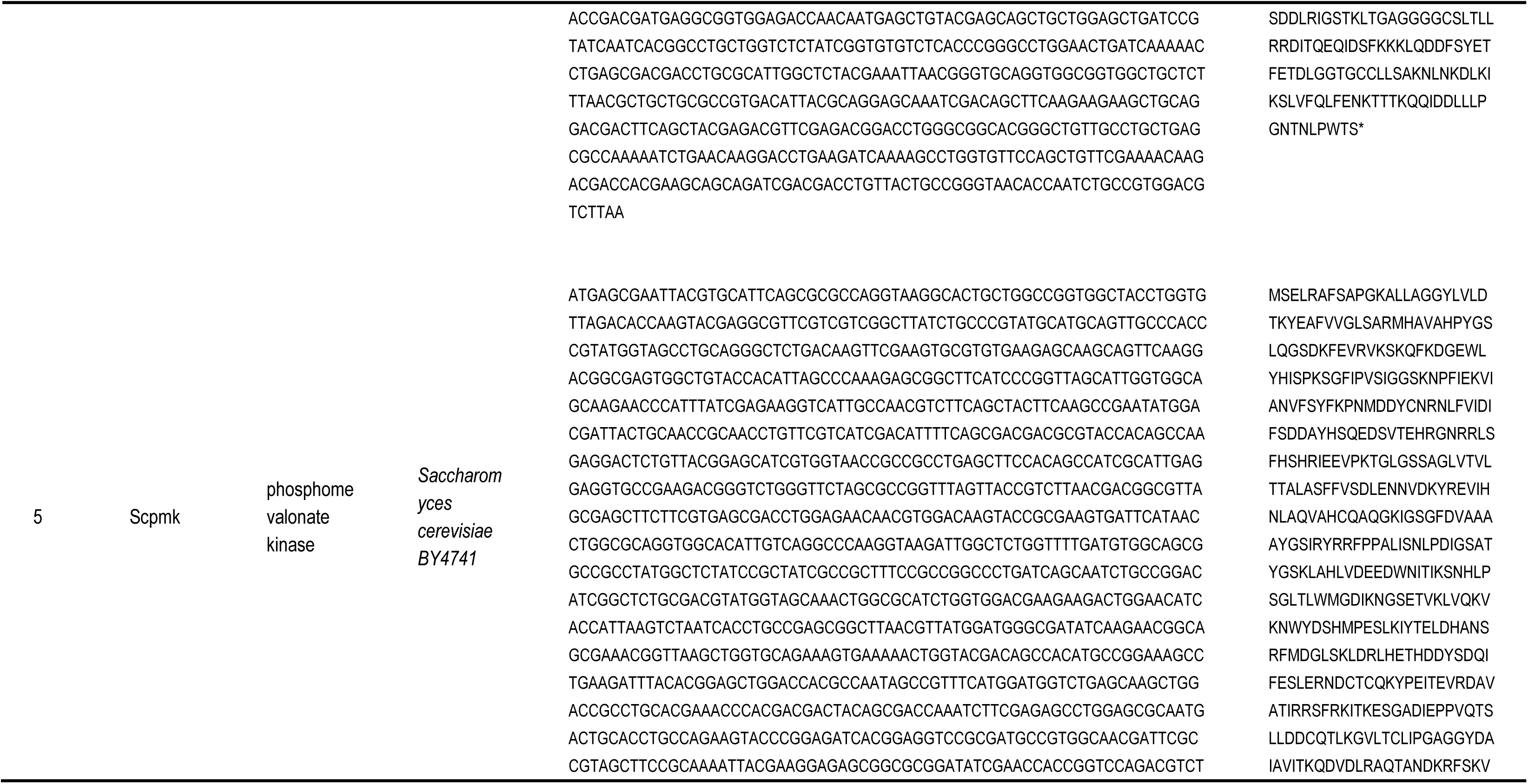
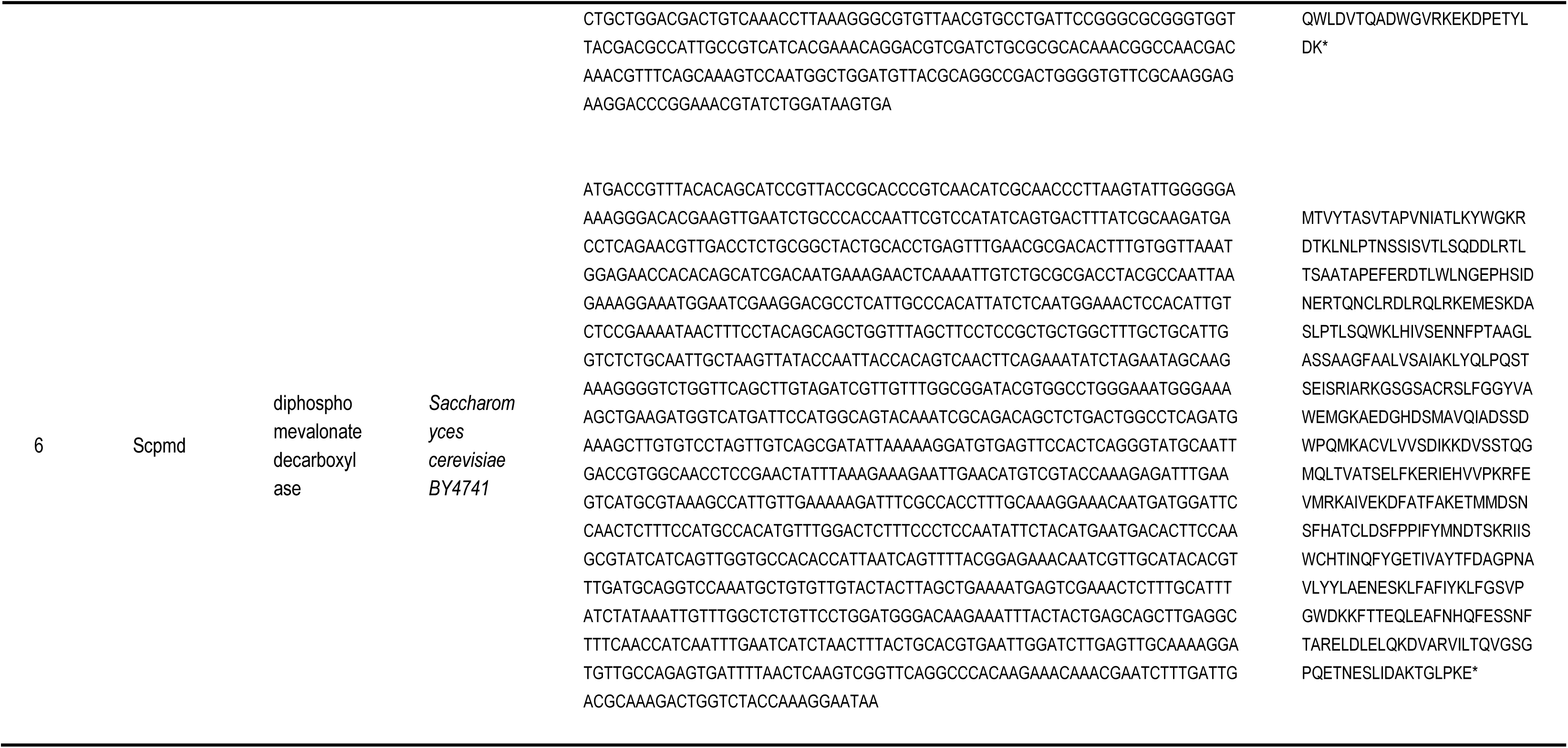
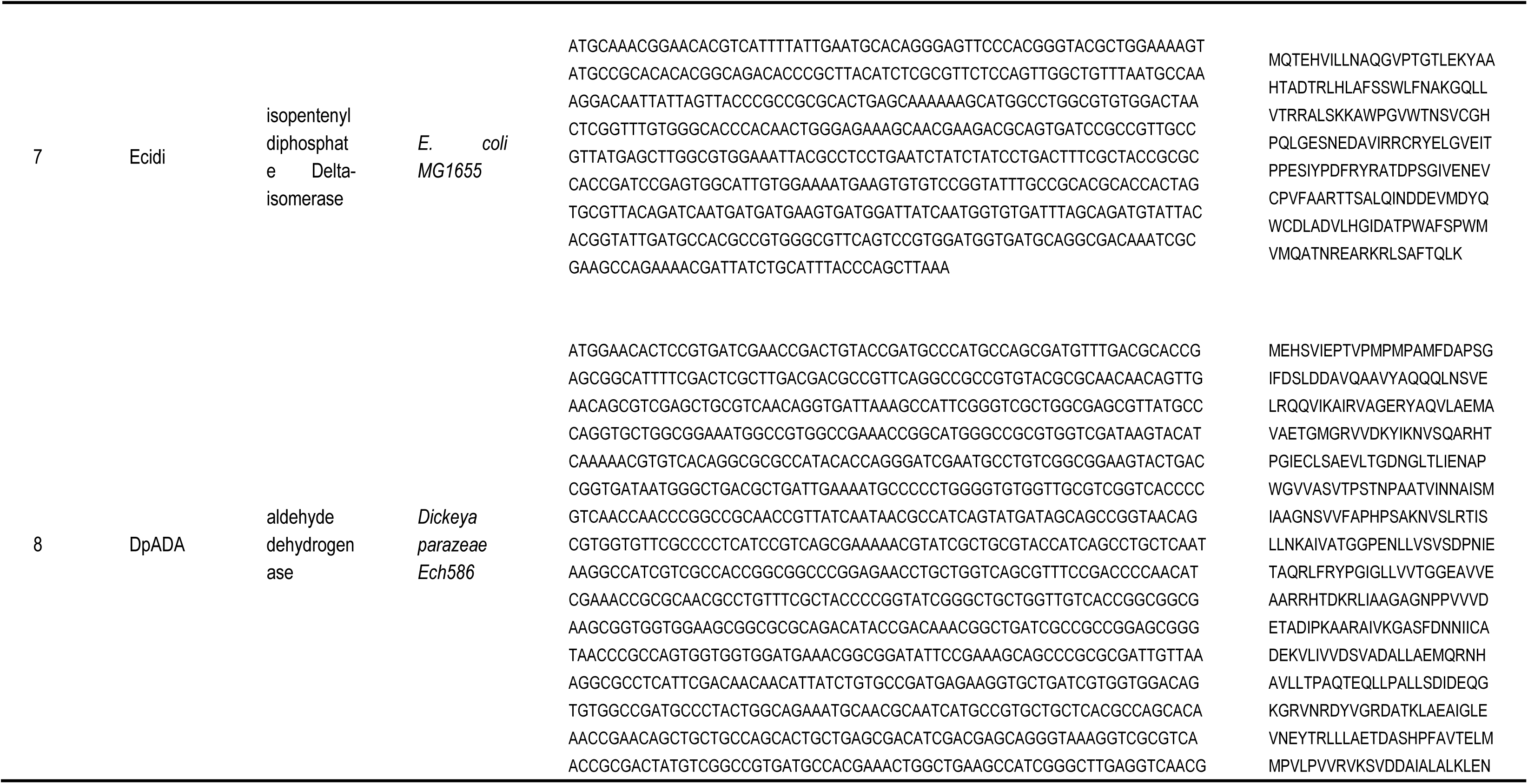
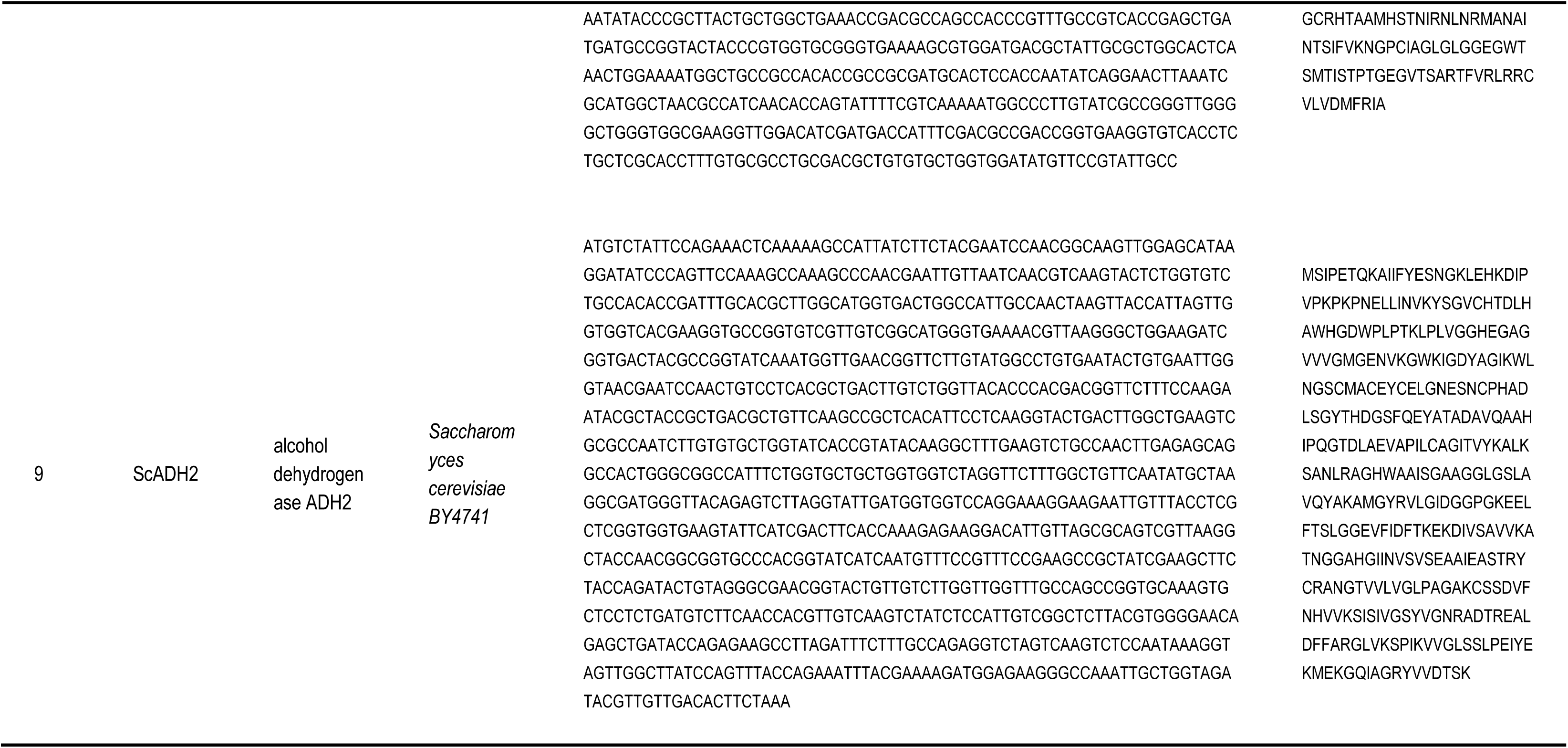
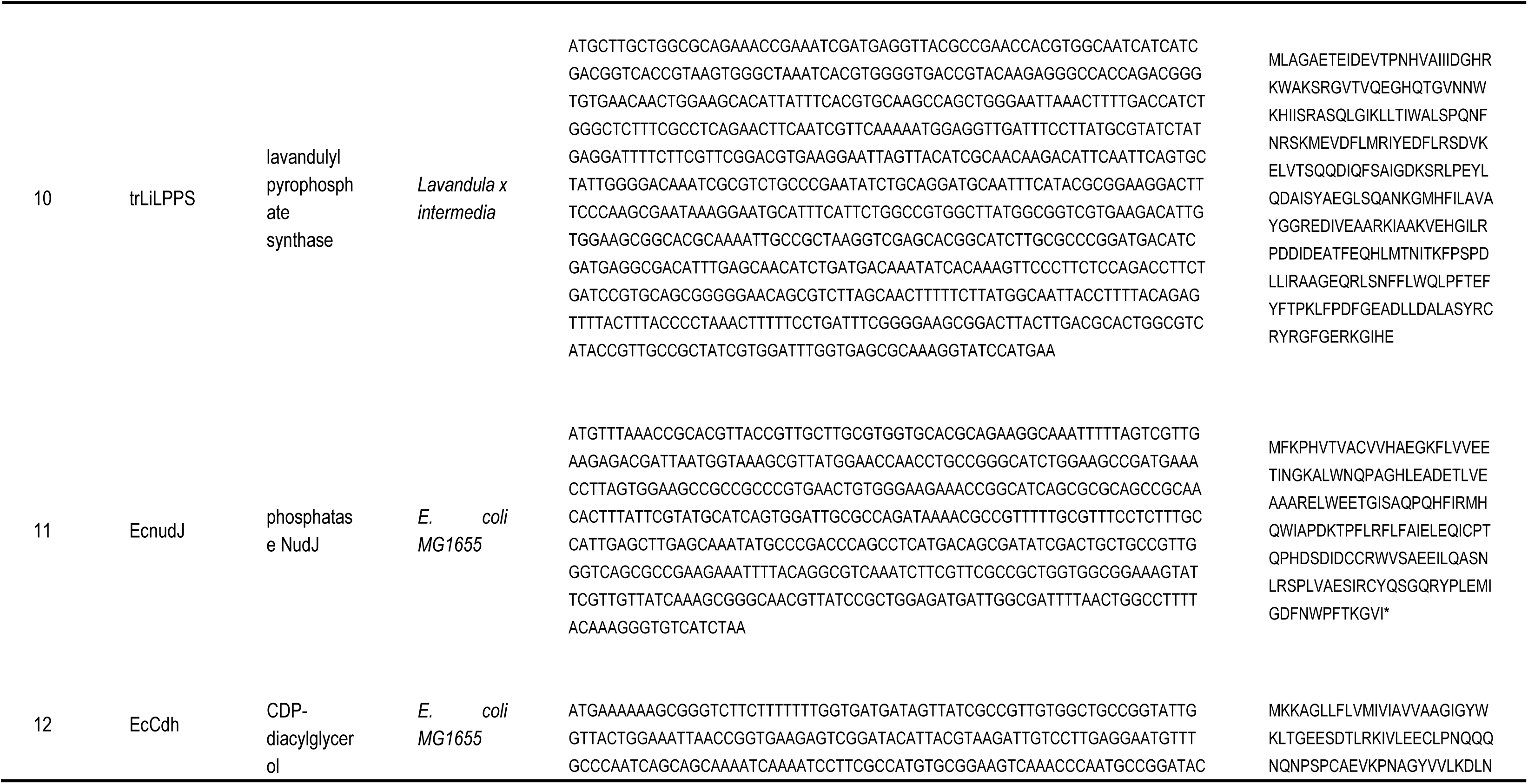
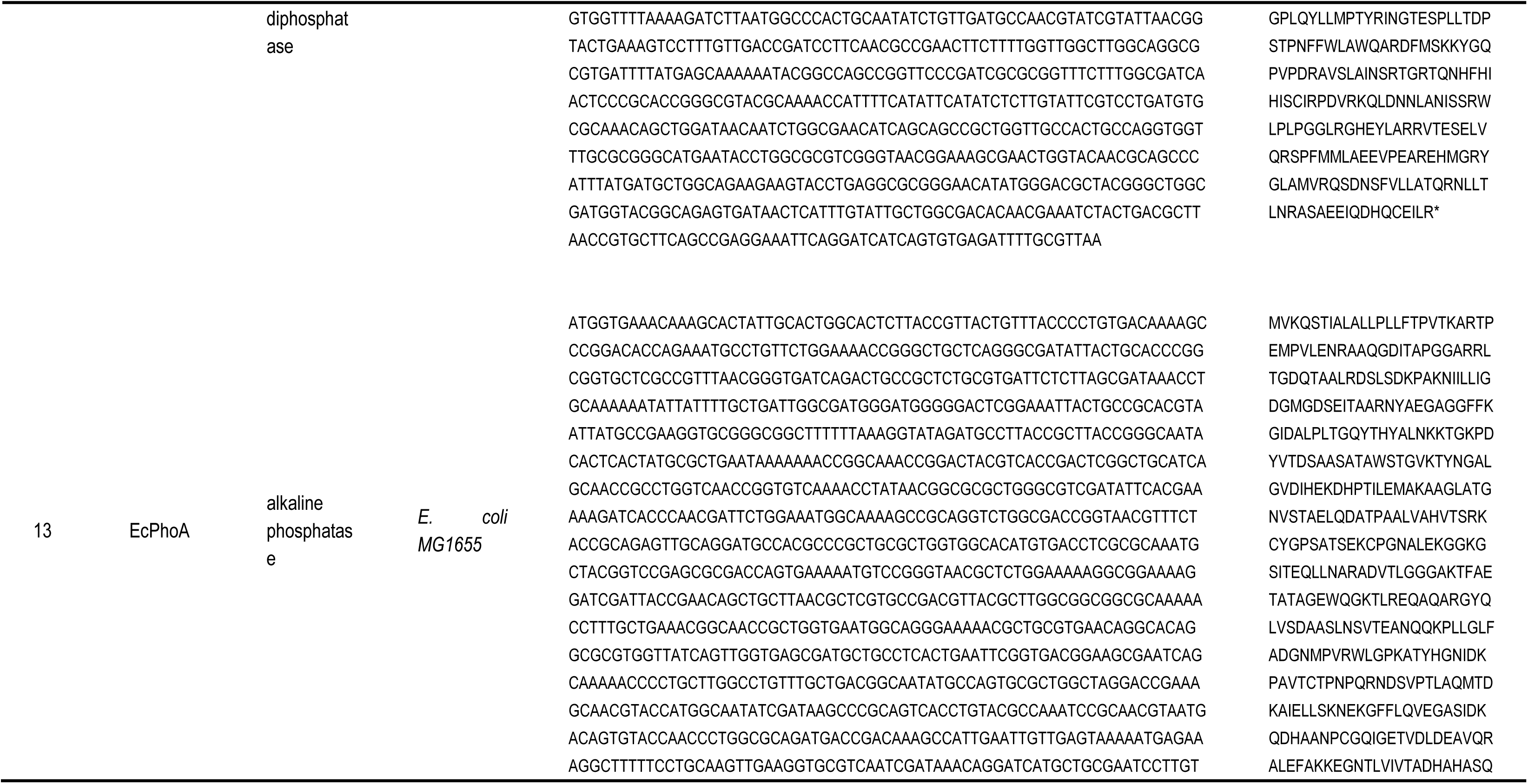
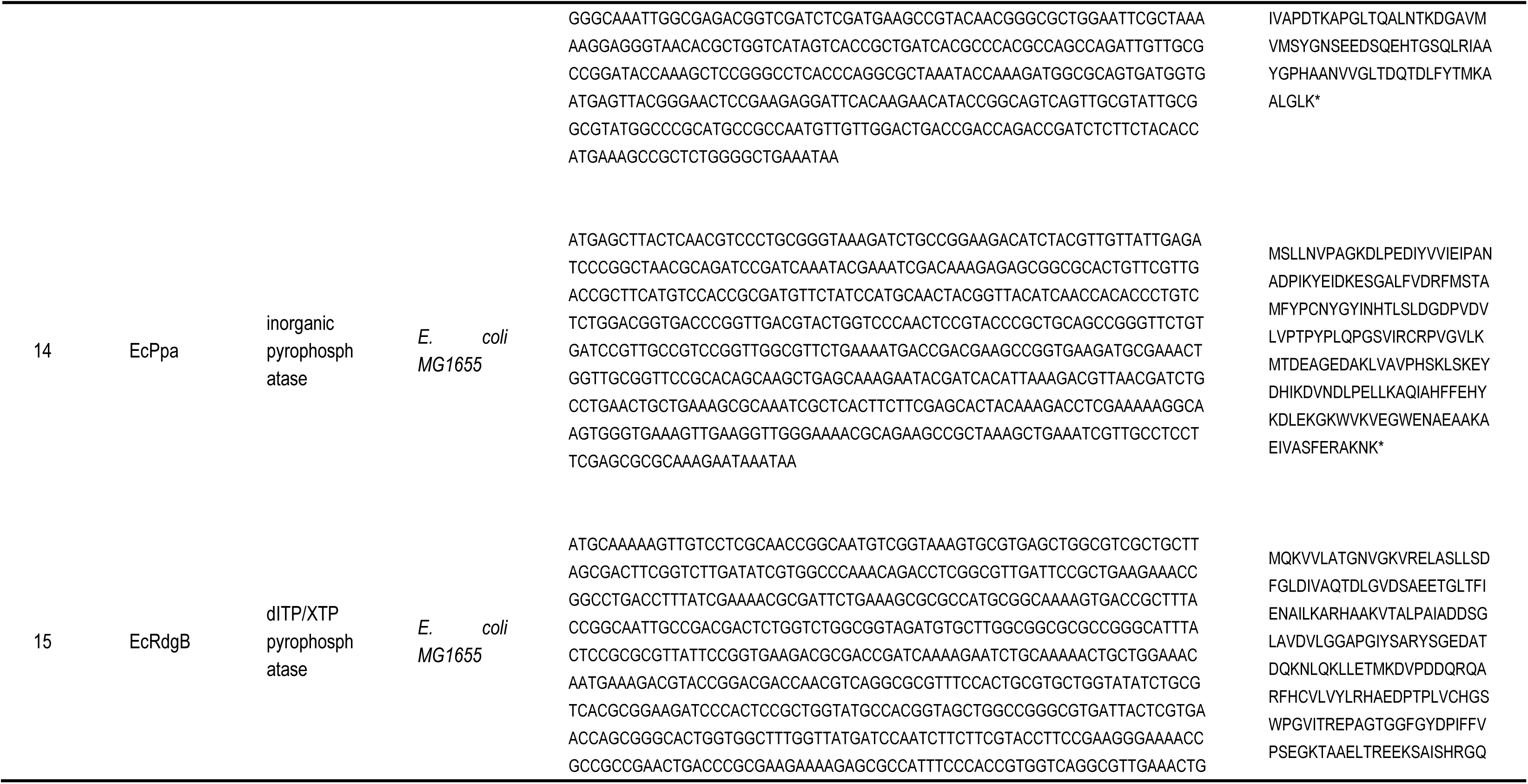
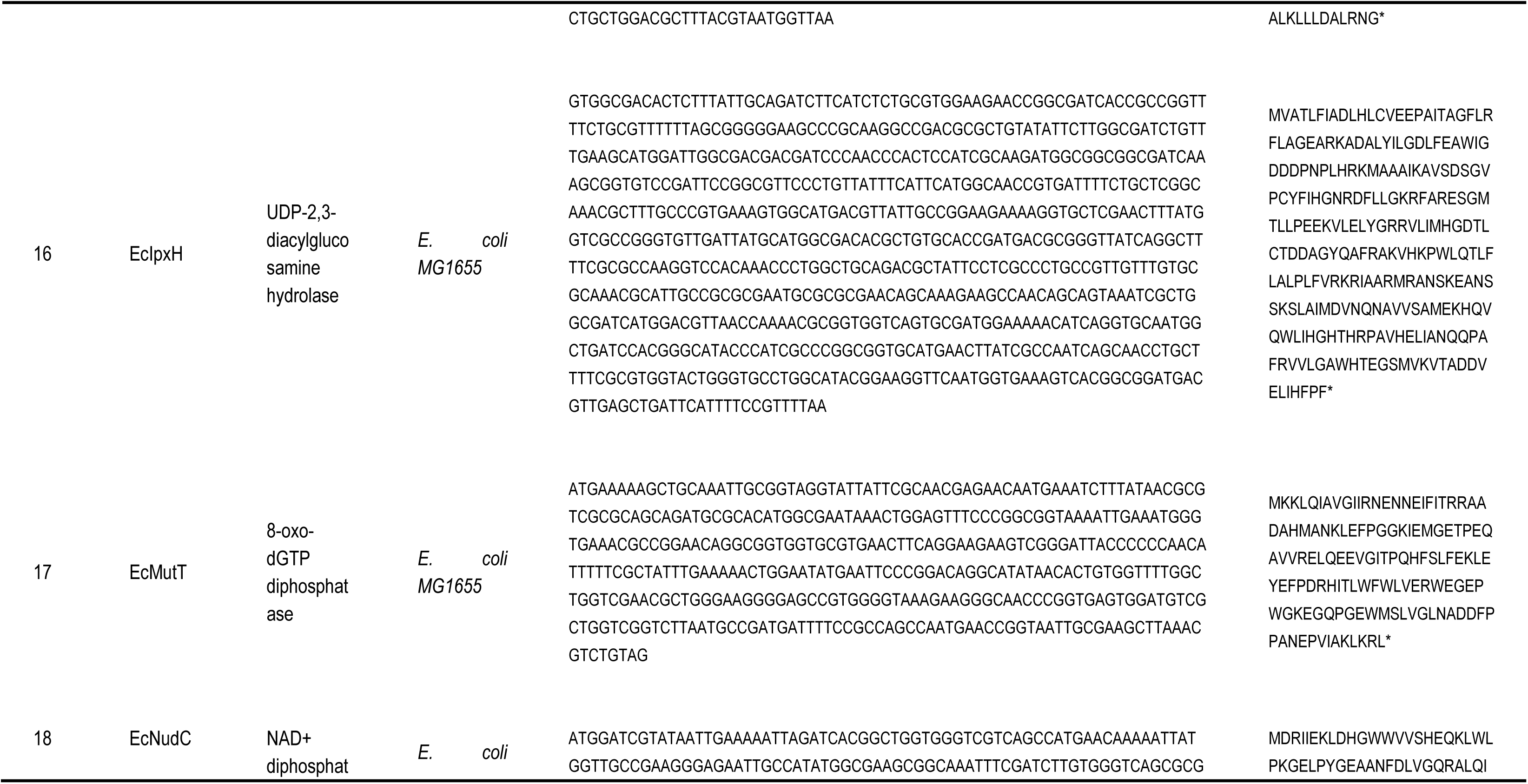
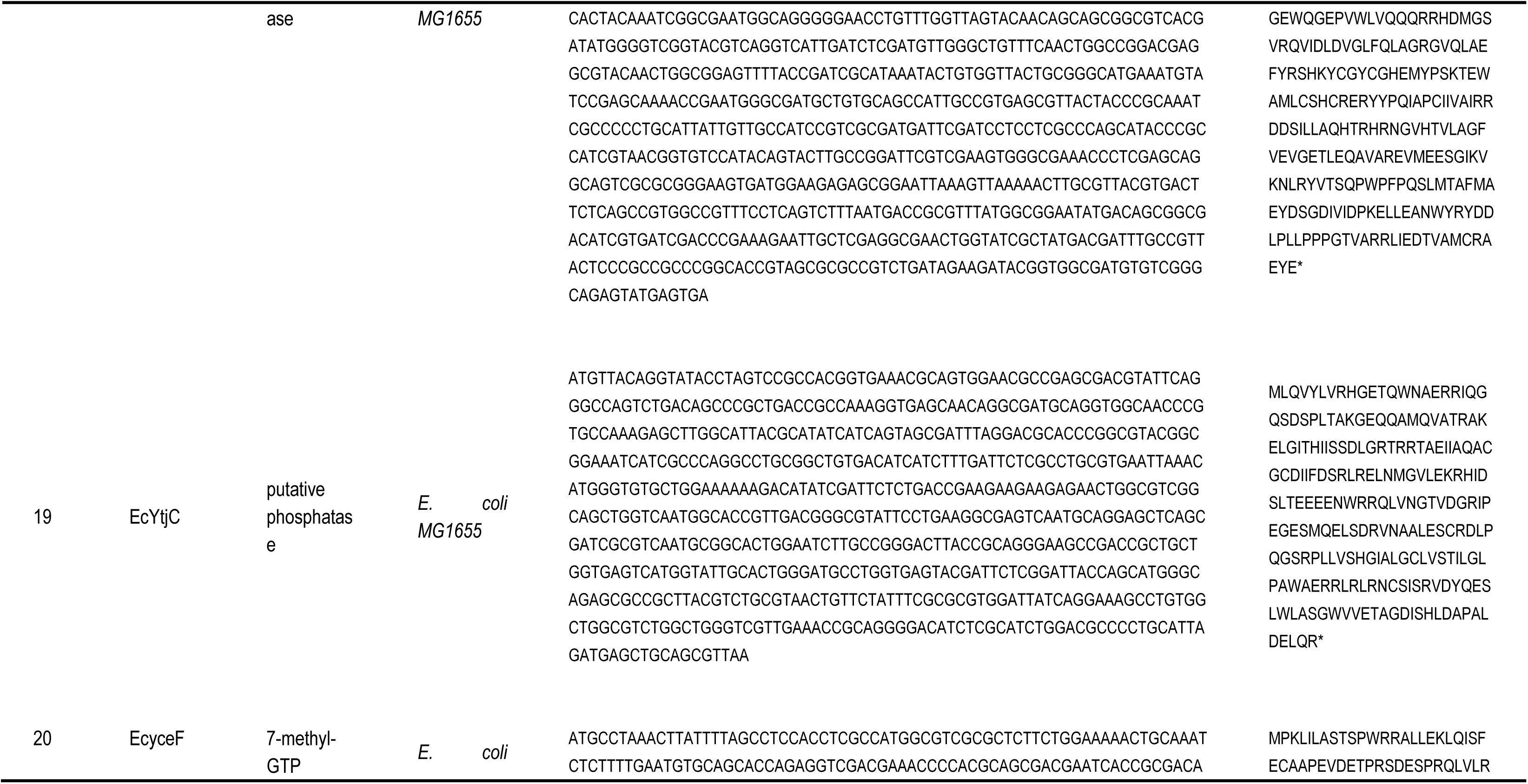
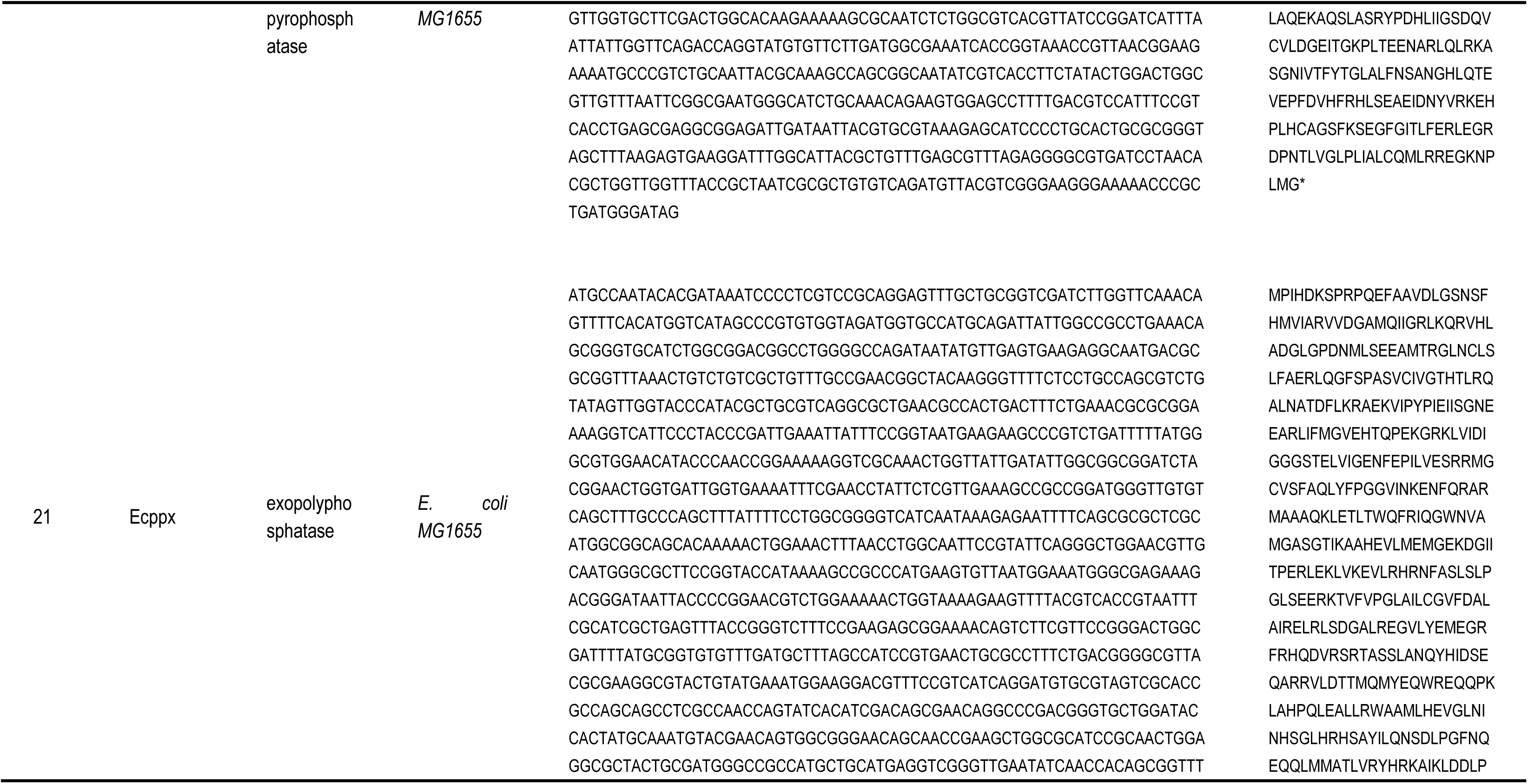
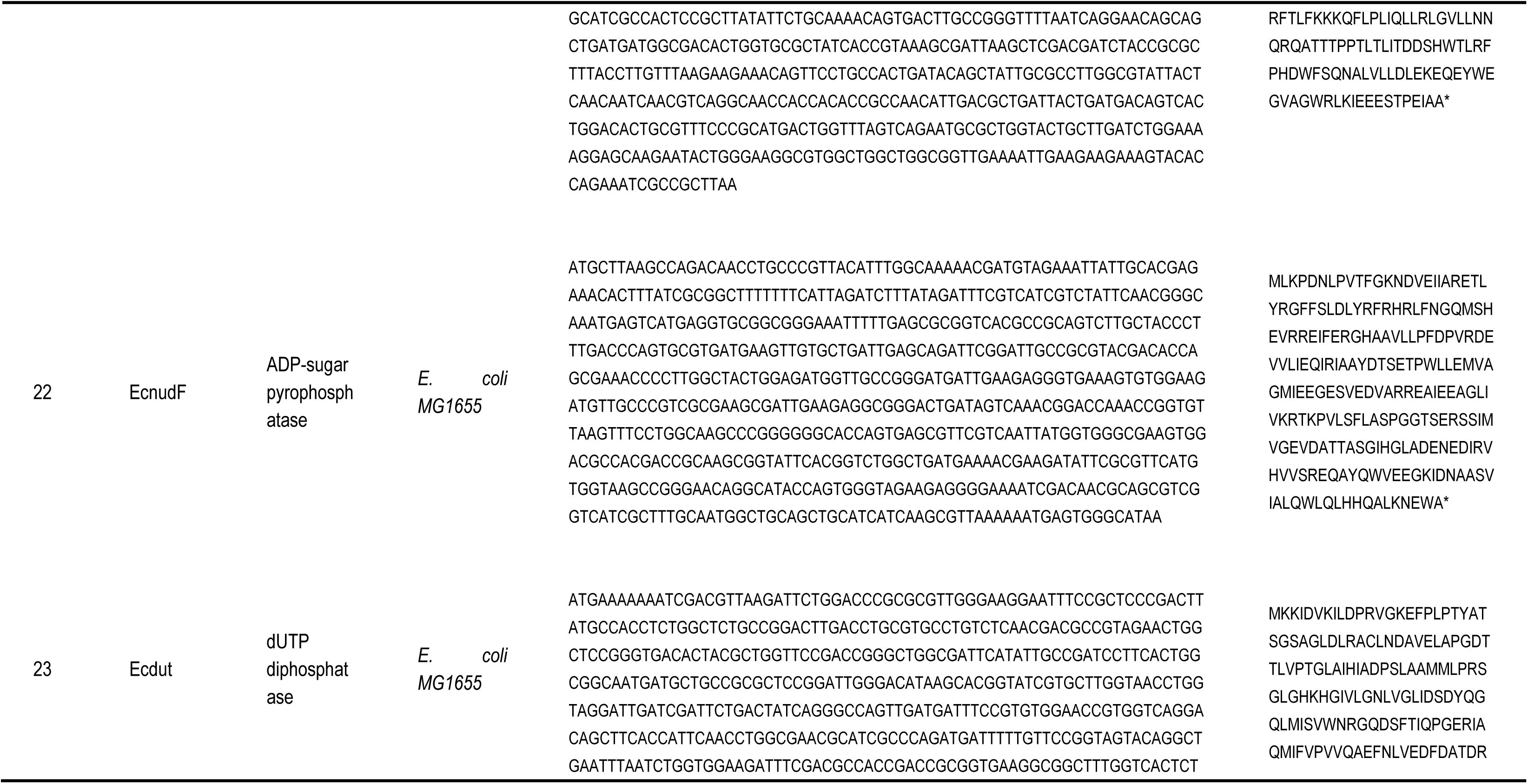
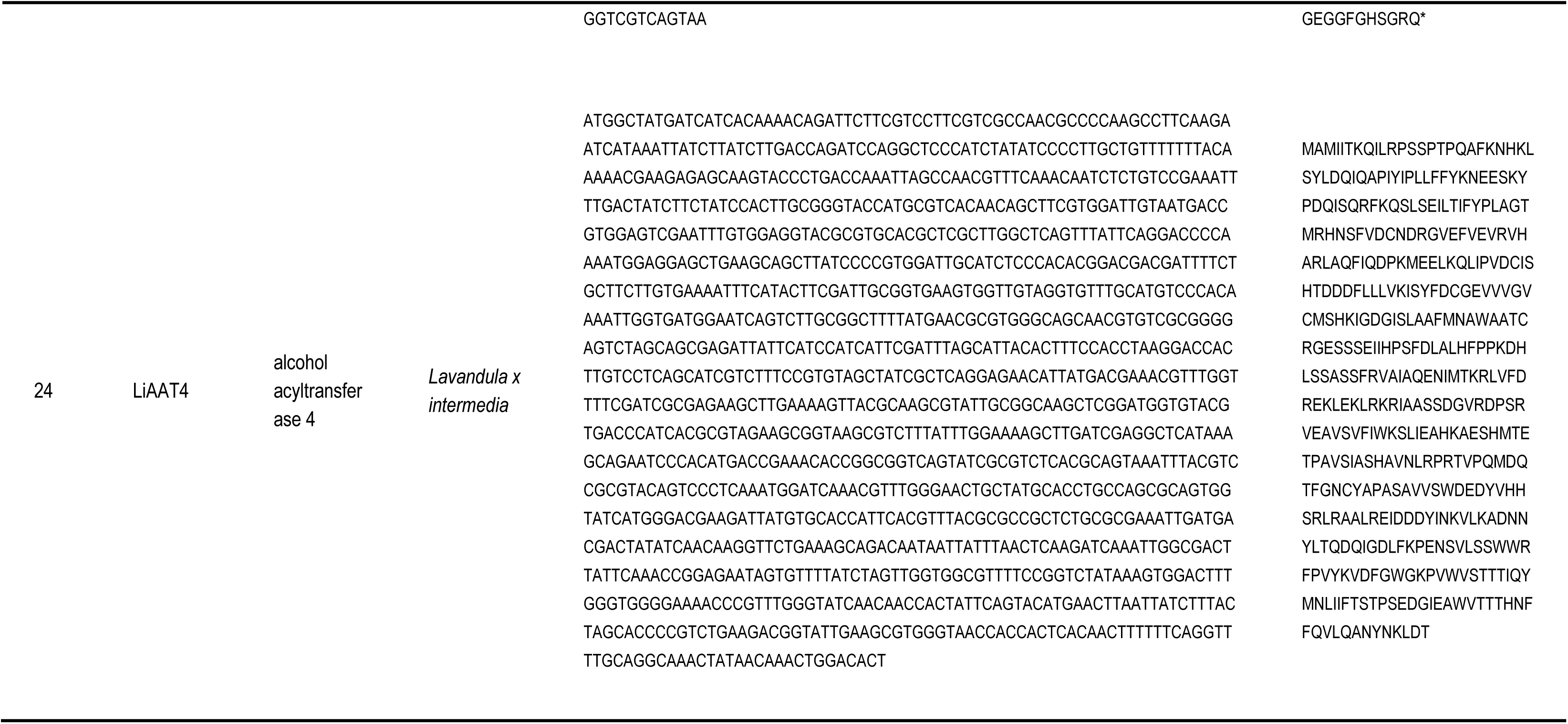
Gene and protein sequence.

### 2.3 Strain cultivation

A single colony of **Lav1-13** was picked from an LB agar plate and inoculated into 2 mL of LB medium supplemented with 50 µg/mL spectinomycin and 50 µg/mL carbenicillin (for lavandulyl acetate production, **LavanA** was cultured in LB medium additionally containing 50 µg/mL kanamycin). The culture was incubated overnight at 37 °C with shaking at 250 rpm. The overnight seed culture was then inoculated at 2.0 % (v/v) into 2 mL of M2 medium containing the appropriate antibiotics and grown at 37 °C, 250 rpm until the optical density at 600 nm (OD600) reached 0.8∼1.0. Protein expression was induced by adding isopropyl β-D-1-thiogalactopyranoside (IPTG, 0.1 mM) and 3-oxohexanoyl-homoserine lactone (3OC6-HSL, 0.1 μM) at specified concentrations (for lavandulyl acetate production, 0.1 mM cumic acid added additionally). Additionally, hexadecane (20%, v/v) and ethanol (final concentration was 10 g/L) were added as an overlay for in situ product extraction. The induced culture was further incubated at 30 °C and 250 rpm for 48 h. After cultivation, the broth was centrifuged at 13,000 rpm for 2 min. For Gas Chromatography-Mass Spectrometry (GC-MS) analysis, 10 µL of the upper hexadecane phase was thoroughly mixed with 90 µL of ethyl acetate before injection.

### 2.4 Analytical methods

The GC-MS analysis was performed using a system of 7890B-5977B (Agilent Technologies, Santa Clara, CA, USA) equipped with HP-5MS Capillary Column (30 m × 0.25 mm × 0.25 μm, Agilent Technologies, CA, USA). After fermentation, the mixture was collected and centrifuged at 13,000 rpm for 10 min. One microliter of supernatant was injected into the GC in split 1:10 mode. The oven temperature was programmed to be maintained at the initial temperature of 50 °C for 1 min and then increased to 100 °C at a rate of 5 °C/min, held for 1 min, increased to 200 °C at 5 °C/min, held for 1 min, increased to 300 °C at 5 °C/min, held for 5 min. The temperatures of the MS source and MS quad were set at 230 °C and 150 °C, respectively. The scan range width was 40–500 m/z.

## 3. Results and Discussion

### 3.1. Screening PPase to biosynthesize lavandulol from glycerol

Although the exact phosphatase directly responsible for catalyzing the conversion of lavandulyl diphosphate (LPP) to lavandulol remains to be fully characterized, it is well recognized that LPP serves as the immediate biosynthetic precursor of lavandulol. Given the structural similarity between LPP and known nucleotide substrates, we hypothesized that endogenous PPase in *E. coli* might possess latent promiscuous activity toward LPP via nonspecific diphosphate hydrolysis (**Fig. 1**). To systematically identify such catalytic candidates, we selected 13 PPases from *E. coli*, including well-characterized members of the Nudix hydrolase superfamily (e.g., NudF, RdgB, MutT), as well as other phosphatases with potential pyrophosphatase activity. Each candidate gene was individually cloned into a modular plasmid backbone (**pLav1-13+EUP**, AmpR+pMB1) co-expressing the genes coding the enzymes in EUP and the downstream trLiLPPS[13] responsible for LPP biosynthesis. This construct was introduced into *E. coli* harboring **pMVA**, a plasmid encoding the MVA pathway enzymes for IPP/DMAPP supply[14], thereby generating the screening strain **Lav1-13**.

**Fig. 1.**
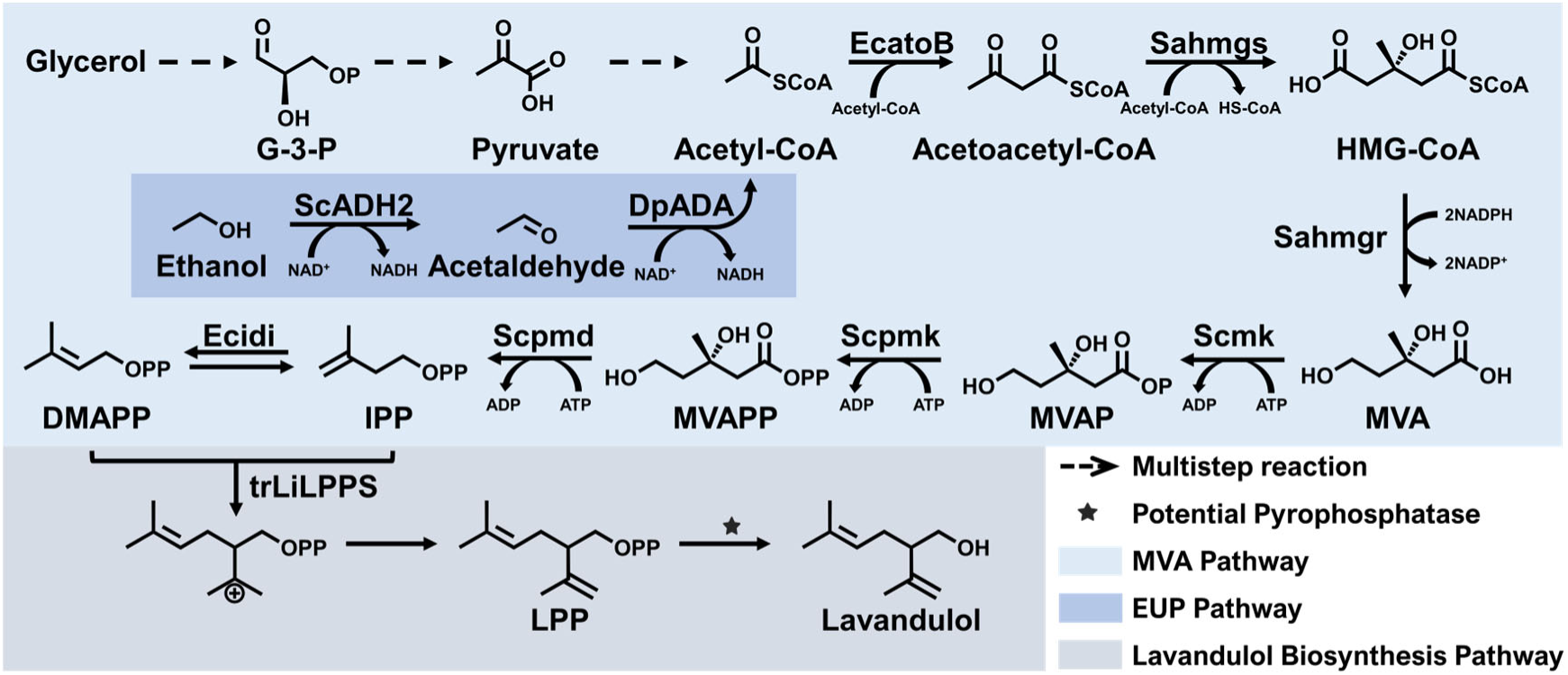
Biosynthetic pathway of lavandulol produced from glyceryl in *E. coli*. G-3-P: glyceraldehyde 3-phosphate; HMG-CoA: 4-hydroxy-3-methyl-glutaryl-CoA; MVAP: mevalonate 5-phosphate; MVAPP: mevalonate 5-diphosphate; IPP: isopentenyl pyrophosphate; DMAPP: dimethylallyl pyrophosphate; GPP: geranyl pyrophosphate; EcatoB: acetyl-CoA acetyltransferase; Schmgs: hydroxymethylglutaryl-CoA synthase; Schmgr: hydroxymethylglutaryl-CoA reductase; Scmk: mevalonate kinase; Scpmk: phosphomevalonate kinase; Scpmd: diphosphomevalonate decarboxylase; Ecidi: isopentenyl-diphosphate Delta-isomerase; ScADH2: alcohol dehydrogenase ADH2; DpADA: aldehyde dehydrogenase; trLiLPPS: truncated lavandulyl pyrophosphate synthase. The details of the genes overexpressed in the engineered *E. coli* in this study can be found in **Table 3**.

Fermentation were performed under the induction and hexadecane overlay conditions to enable *in situ* extraction of the volatile lavandulol. GC-MS analysis revealed that among all tested PPase, RdgB exhibited the highest activity, yielding 24.9 mg/L of lavandulol, followed by NudF, which achieved 14.9 mg/L (**Fig. 2a**). Notably, most other candidates displayed negligible or undetectable activity, suggesting that dephosphorylation of LPP is a highly selective process requiring specific structural complementarity. These results were further validated by comparative mass spectrum quantification and retention time alignment with an authentic lavandulol standard (**Fig. 2b**). EcRdgB was reported as a protein with deoxyribonucleoside triphosphate pyrophosphohydrolase (dNTPase) activity, specifically acting on non-canonical DNA precursors like dITP and dXTP[15]. The superior catalytic efficiency of RdgB toward LPP underscores its promiscuity of substrate flexibility and highlights its potential as a non-natural monoterpene phosphatase. Importantly, this work represents the demonstration of lavandulol biosynthesis in *E. coli* via a rationally designed LPP dephosphorylation module. Based on these findings, RdgB was selected as the core enzyme for subsequent pathway engineering, laying a biochemical foundation for the *de novo* microbial biosynthesis of lavandulol and its derivatives.

**Fig. 2.**
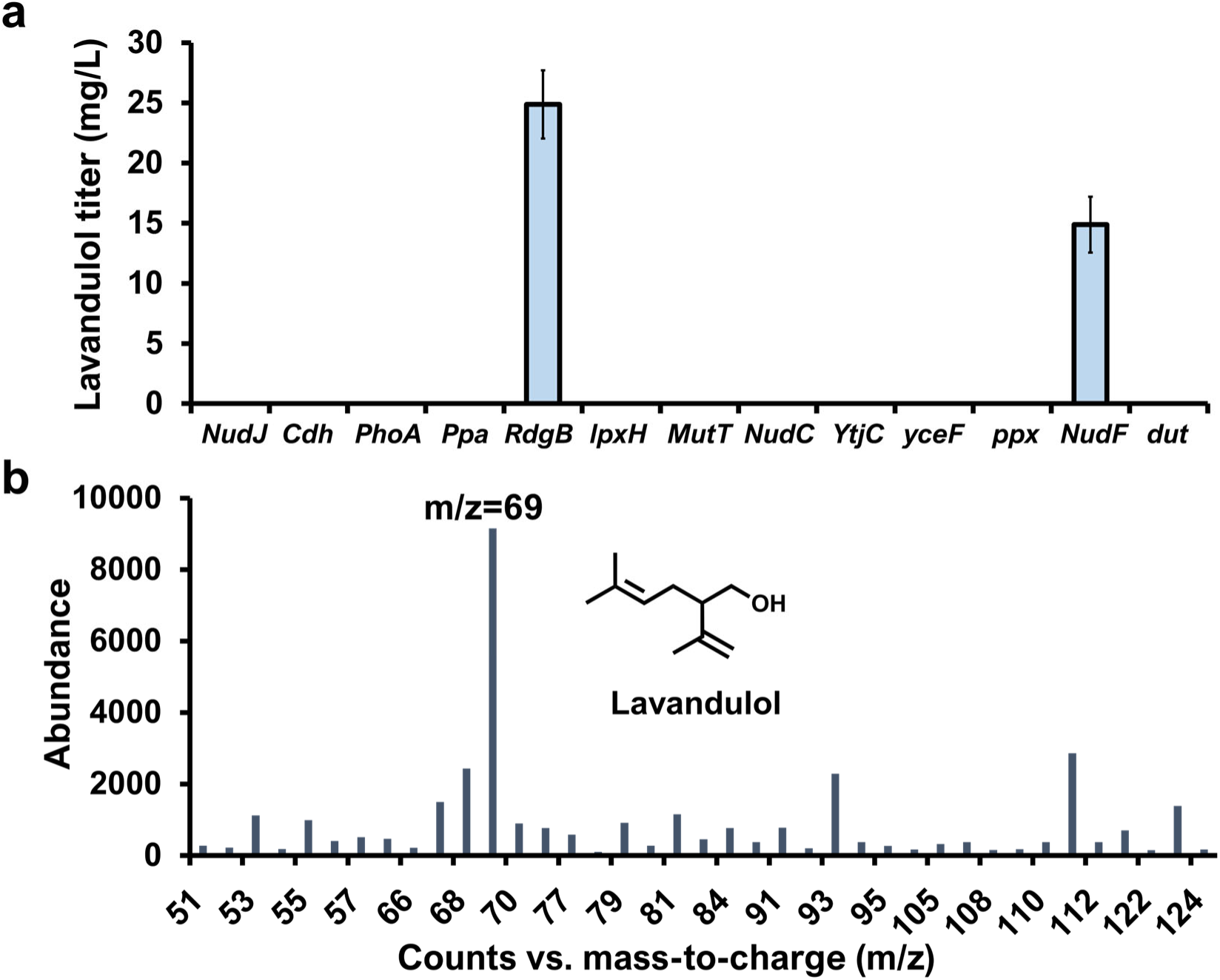
Screening of 13 *E. coli* PPases for lavandulol biosynthesis. (**a**) The results of screening with 13 DTPPs; (**b**) GC-MS mass spectrum of lavandulol produced by **Lav5.**

Interestingly, our study also highlights the previously underexplored role of endogenous PPases in modulating isoprenoid biosynthesis in engineered *E. coli*. The ability of RdgB and NudF to catalyze the hydrolysis of lavandulyl diphosphate suggests that endogenous PPases may possess broad substrate flexibility toward isoprenoid pyrophosphates. This enzymatic promiscuity opens new avenues for expanding microbial production of diverse isoprenoid derivatives. In particular, tailored PPases could be exploited to selectively hydrolyze C5 (e.g., IPP, DMAPP), C10 (e.g., GPP, LPP), or C15 (e.g., FPP) precursors, enabling the development of novel biosynthetic routes for the generation of structurally varied C5-C15 isoprenoid alcohols or related molecules. Rational mining and engineering of such PPases thus represent a promising direction for broadening the enzymatic toolbox in synthetic isoprenoid biology. Recently, Nie et al.[9] demonstrated the feasibility of *de novo* lavandulol biosynthesis in *S. cerevisiae*. However, this study primarily focused on the expression and metabolic optimization of trLiLPPS without using any dephosphorylation PPase to form lavandulol. Since *in vitro* enzymatic assay using purified LiLPPS with DMAPP as the substrate required additional hydrolysis by calf intestinal alkaline phosphatase to release lavandulol[13], it remains unclear whether endogenous PPases in *S. cerevisiae* were responsible for LPP hydrolysis *in vivo*. This uncertainty indicates the need for *in vitro* enzymatic validation to determine whether yeast-derived phosphatases exhibit activity toward irregular monoterpene diphosphates such as LPP.

### 3.2. Lavandulyl acetate biosynthesis in *E. coli*

Following the successful establishment of lavandulol biosynthesis in *E. coli*, we further expanded the biosynthetic scope to include lavandulyl acetate, a commercially valuable ester derivative widely utilized in fragrance and cosmetic industries due to its desirable olfactory properties. The esterification of lavandulol into lavandulyl acetate can be achieved using AAT with acetyl-CoA as acyl donor. To achieve the biosynthesis of lavandulyl acetate, we constructed an engineered *E. coli* strain (**pLiAAT4K**) harboring the LiAAT4 gene from *Lavandula* x *intermedia* encoding AAT, under the control of a cumate-inducible CymR promoter. LiAAT4 was selected based on its reported capability to catalyze the esterification of monoterpene alcohols, including lavandulol[11]. The resultant strain (**pLiAAT4K**, KanR+pCDF) was subsequently transformed into **Lav5** (already engineered for lavandulol production) to generate the strain integrated three-plasmid as **LavanA** (**Fig. 3**).

**Fig. 3.**
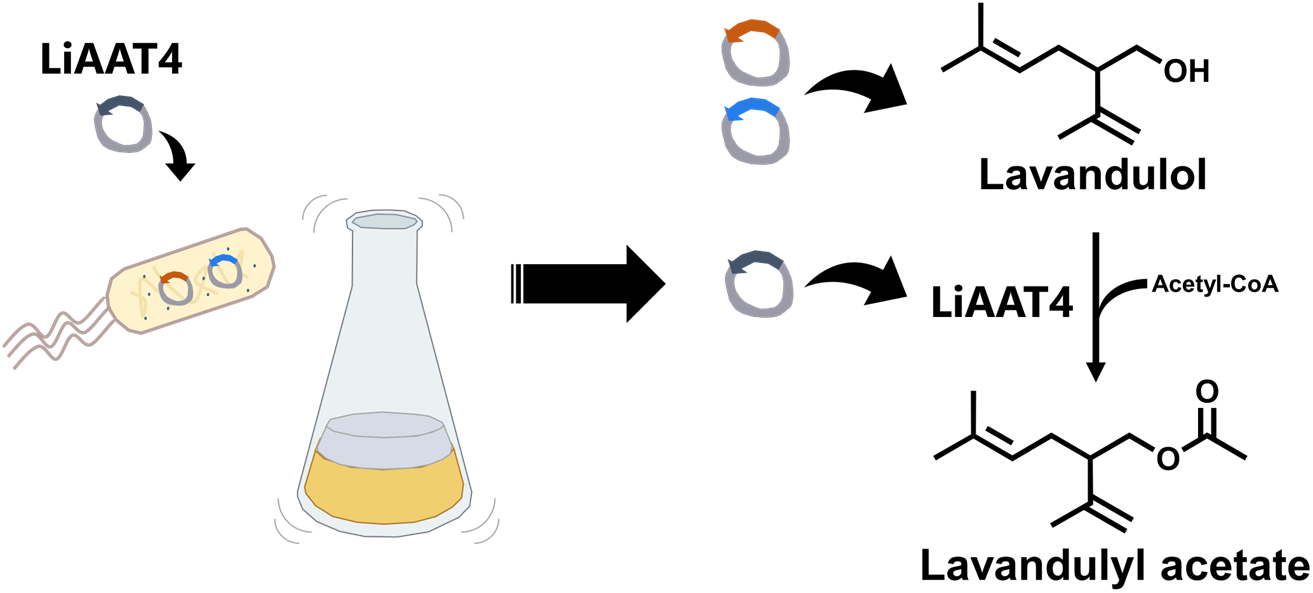
Modular biosynthesis of lavandulyl acetate in engineered *E. coli*. The three-plasmid system includes the MVA pathway, lavandulol biosynthesis module, and cumate-inducible LiAAT4. Lavandulol is converted to lavandulyl acetate through LiAAT4-catalyzed esterification, with acetyl-CoA as the acyl donor.

Under optimized induction conditions (0.1 mM IPTG, 0.1 μM 3OC6-HSL, and 0.1 mM cumic acid), we evaluated the capability of this microbial platform to directly synthesize lavandulyl acetate from glycerol-derived precursors. The fermentation was performed in shake-flasks with a hexadecane overlay to facilitate *in situ* extraction and minimize product loss through volatility and cellular metabolism. Additionally, considering that ethanol is a commonly employed co-substrate for enhancing acetyl-CoA supply, we assessed the effect of ethanol supplementation (10 g/L) on lavandulyl acetate production. The fermentation data revealed a notably higher production titer of lavandulyl acetate (42.4 mg/L) without ethanol addition compared to ethanol-supplemented conditions (**Fig. 4a**). GC-MS analysis further confirmed the successful production of lavandulyl acetate, identified by matching its retention time and mass spectrum to those of a commercially available authentic standard (**Fig. 4b**). The decrease in product titer upon ethanol supplementation was not due to a limitation in acetyl-CoA supply. Instead, the additional ethanol likely exacerbated DMAPP accumulation, which can lead to cytotoxicity and metabolic imbalance, ultimately suppressing overall product formation.

**Fig. 4.**
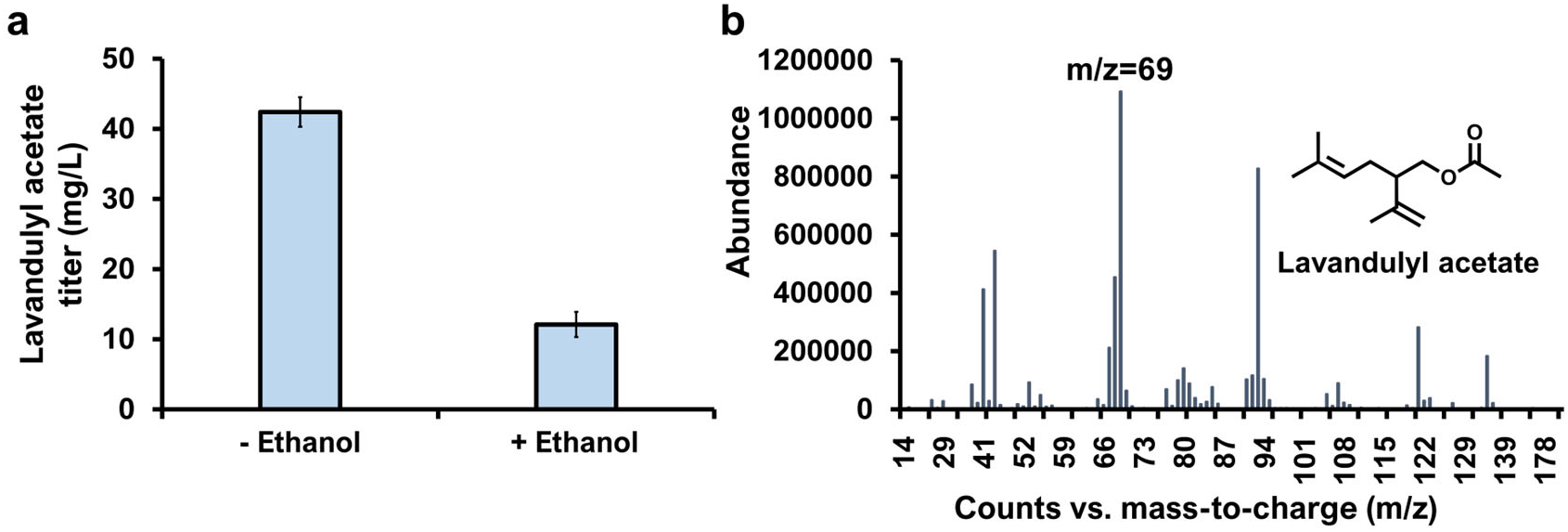
Biosynthesis of lavandulyl acetate from glycerol with or without EUP. (**a**) The result of fermentation of **LavanA** with/without the addition of 10 g/L of ethanol; (**b**) GC-MS mass spectrum of lavandulyl acetate produced by **LavanA.**

These findings demonstrate the production of lavandulyl acetate using an engineered microbial platform, highlighting the feasibility of integrating monoterpene biosynthesis with the AAT-mediated esterification pathways in a single microbial host. Although the current production titer remains relatively low, the established three-plasmid system provides a solid foundation for further optimization. Future efforts may focus on engineering key catalytic steps, such as protein engineering of trLiLPPS and EcRdgB, to enhance their catalytic efficiencies and thereby improve the overall carbon conversion rate of the biosynthetic pathway. Given the observed accumulation bottleneck, it appears that lavandulol biosynthesis remains the primary limiting step, suggesting that boosting lavandulol availability will be crucial for further enhancing overall acetate ester production.

## 4. Conclusion

This study reports the *de novo* microbial production of lavandulol and lavandulyl acetate in *E. coli* using a modular three-plasmid system. Through systematic screening, RdgB was identified as an efficient LPP phosphatase, enabling lavandulol biosynthesis in *E. coli*. Further co-expression of LiAAT4, an AAT from *Lavandula* x *intermedia*, enabled the bioconversion of lavandulol into lavandulyl acetate, reaching titers of 24.9 mg/L and 42.4 mg/L, respectively. While the current yield remains moderate, the pathway modularity and host compatibility provide a solid foundation for future improvement. Key strategies include protein engineering of trLiLPPS and EcRdgB, metabolic flux balancing, and fermentation medium/process optimization. Our work establishes a versatile platform for the biosynthesis of irregular monoterpene and ester derivative, advancing the sustainable production of high-value fragrance compounds.

## 5. Acknowledgments

This work was supported by the National Key R&D Program of China (2022YFD2101304), the Start-up fund of Shanghai Jiao Tong University (WH220415004), the Shanghai Collaborative Innovation Center of Agri-Seeds (ZXWH3150101/002). We also thank Jie Xu and Qian Luo for the guidance of using GC-MS in Public Instrument Service Platform of School of Life Sciences, Shanghai Jiao Tong University.

## 6. Author Contributions

Dianqi Yang: Data curation, Methodology, Investigation, Formal analysis, Resources, Validation, Writing-Original Draft. Xiaoqiang Ma: Conceptualization, Data curation, Supervision, Project administration, Funding acquisition, Formal analysis, Visualization, Writing-Original Draft, Writing - Review & Editing.

## 7. Declarations

The plasmid of pMVA and the flow used in the strain producing geraniol has been disclosed in a patent submitted: Ma, X. & Yang, D. A method for biosynthesizing (R)-lavandulol and (R)-lavandulyl acetate. Publication Number: CN116926130A (Shanghai Jiao Tong University, China).

